# Sensorimotor temporal recalibration: the contribution of motor-sensory and inter-sensory components

**DOI:** 10.1101/2021.03.26.437189

**Authors:** Belkis Ezgi Arikan, Bianca M. van Kemenade, Katja Fiehler, Tilo Kircher, Knut Drewing, Benjamin Straube

**Affiliations:** Experimental Psychology, Justus Liebig University Giessen, Otto-Behaghel Str. 10F, D-35394 Giessen, Germany; Institute of Neuroscience and Psychology, University of Glasgow, 58 Hillhead Street, G12 8QB, Glasgow, UK; Department of Psychiatry and Psychotherapy, Philipps University Marburg, Rudolf-Bultmann-Str. 9, 35039, Marburg, Germany

**Keywords:** adaptation, auditory, visual, cross-modal transfer, efference copy

## Abstract

Adaptation to delays between actions and sensory feedback is important for efficiently interacting with our environment. Adaptation may rely on predictions of action-feedback pairing (motor-sensory component), or predictions of tactile-proprioceptive sensation from the action and sensory feedback of the action (inter-sensory component). Reliability of temporal information might differ across sensory feedback modalities (e.g. auditory or visual), influencing adaptation. Here, we investigated the role of motor-sensory and inter-sensory components on sensorimotor temporal recalibration for motor-auditory events (button press-tone) and motor-visual events (button press-Gabor patch). In the adaptation phase of the experiment, the motor action-feedback event pairs were presented with systematic temporal delays (0ms or 150ms). In the subsequent test phase, sensory feedback of the action were presented with variable delays. The participants were then asked whether this delay could be detected. To disentangle motor-sensory from inter-sensory component, we varied movements (active button press or passive depression of button) at adaptation and test. Our results suggest that motor-auditory recalibration is mainly driven by motor-sensory component, whereas motor-visual recalibration is mainly driven by inter-sensory component. Recalibration transferred from vision to audition, but not from audition to vision. These results indicate that motor-sensory and inter-sensory components of recalibration are weighted in a modality-dependent manner.

## Introduction

Perceiving sensory events almost always involves dealing with temporal discrepancies. Discrepancies may result from temporal differences in neural transduction ^1,2^, developmental changes ^3^, or physical characteristics of a given sensory input ^4^. Yet, humans are highly efficient in compensating for these discrepancies. For example, temporal misalignment between our actions and their sensory consequences resulting from any or a combination of the above-mentioned factors can be accommodated. This is known as sensorimotor temporal recalibration, which has been demonstrated for actions leading to a specific sensory consequence ^5–10^ as well as for actions leading to multiple sensory consequences ^11–13^. Sensorimotor temporal recalibration aids not only in the binding of sensory and motor events that belong together, but also in attributing control (agency) over the events we generate ^14^. This suggests that temporal recalibration is specific to the events that are causally-related. However, voluntary actions seem to provide us with an additional advantage in recalibrating temporal perception ^15^. Actions can trigger an internal forward model which predicts the sensory consequences of the action based on the efference copy of the motor command ^16–18^. Such predictive processing may lead to stronger adaptation of motor-sensory pairings relative to purely sensory event pairings. Indeed, it has been consistently demonstrated that actions provide a temporal window into which binding of sensory events can be facilitated ^12,15,19–21^.

Temporal discrepancies between actions and sensory feedback result in misalignment of two components: a motor-sensory component (misalignment between the action and its sensory feedback), and an inter-sensory component (misalignment between crossmodal sensory inputs); one or both needs to be recalibrated ^9,22^. For example, whenever we click a link to a website, some amount of time is required to load and display the page on our computer screen. The *perceived* interval between these events may be due to temporal misalignment between the click and the appearance of the website (motor-sensory component), or between the tactile-proprioceptive feedback arising from the click and the appearance of the website (inter-sensory component). In order to attribute causality between the click and the appearance of the website, the perceived timing of these events are adjusted.

How do motor-sensory and inter-sensory components contribute to sensorimotor temporal recalibration? Research on the dynamics of sensorimotor recalibration has provided insights into this question. In these studies, a sensorimotor event is learned, after which a spatial or temporal perturbation between the events is introduced. The perturbation is modelled as error variance, which can either be random or systematic ^23,24^. Adaptation occurs for systematic errors, which are weighted according to their reliability; i.e., the more reliable estimate with minimum variance receives a higher weight ^23,25–27^. Drawing from this previous work and from work on inter-sensory recalibration ^28,29^, we argue that sensorimotor temporal recalibration results from an optimal combination of motor-sensory and inter-sensory components based on their reliability in minimizing action-feedback discrepancies ^23,25,26,30^. We define temporal recalibration as remapping of the systematic delay to the time point an action-effect pair is expected, leading to the perception of synchrony for the action-feedback event pair. This time point might be an integrated weighted average of time point estimates from motor-sensory and inter-sensory predictions, and the reliability (which is equal to the inverse variance of the time point estimates) of these two types of predictions determines their relative weight:

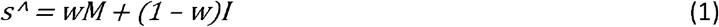

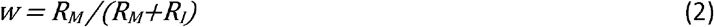

Where *s*^ represents the weighted sum of the individual estimates, *M* represents the motor-sensory component with weight *w*, and *I* represents the inter-sensory component with weight *(1 - w)*. Here, the weight of each component is proportional to the inverse of its variance, *R* – relative to the sum of the inverse variances of the other components. Note that adaptation that is based on more reliable estimates should yield more pronounced adaptation results (or predictions) in short time ^23^, which are weighted higher when different predictions are combined (equations (1) & (2)).

Apart from dynamic modulation of sensorimotor temporal perturbations^24,31^, other studies have focused on the extent to which motor-sensory and inter-sensory components result in adaptation. One approach involves disentangling predictions based on efference copy from reafferent feedback, and testing which component accounts mostly for recalibration. Comparing adaptation during voluntary button presses with a *passive* condition in which the button moved the finger, Stetson et al. ^8^ found larger temporal recalibration effects for voluntary button presses triggering flashes, and smaller recalibration effects for the passive button presses, indicating the importance of action intention (in which efference copy is present) on sensorimotor recalibration. In another study, Arnold et al. ^22^ examined the role of action intentions on sensorimotor temporal recalibration using ballistic reaches with short or longer extent before a voluntary button press triggered a tone. By manipulating the time between the intention to act and the auditory consequence of the action, the authors were able to investigate whether the intention to act or the sensation of having acted drives sensorimotor temporal recalibration. They found that the temporal relationship between tactile signals associated with completion of the action and the auditory feedback of the action determines recalibration rather than the action itself, highlighting the role of the inter-sensory component over the motor-sensory component.

Apart from contributions of motor-sensory and inter-sensory components, the sensory modality of the action feedback (e.g., auditory or visual ^8,10,22^) may also impact temporal recalibration ^10,32^. In general, audition has superior temporal resolution compared to the other senses ^33,34^. This has also been observed when sensory modalities are processed together. For example, audio-tactile events have been found to have higher temporal resolution than visuo-tactile or audio-visual events ^35^, suggesting higher reliability for auditory events in the temporal domain. If this is the case, the optimal combination model should take into account the reliability of the sensory feedback of the action; that is, its variance in remapping the timing of action-feedback events. Sensory feedback reliability might also interact with the reliability of the motor-sensory and inter-sensory components in recalibrating action-feedback timing. One can test this by comparing adaptation effects for learned action-feedback pairs with a condition in which the sensory modality is changed after adaptation ^6,9^. Transfer of recalibration to a different sensory modality (e.g. motor-auditory to motor-visual, or vice versa) indicates a supramodal (modality a-specific) mechanism which is not affected by the sensory feedback associated with the action, whereas absence of transfer suggests a modality-specific mechanism that is influenced by the sensory modality of the action feedback ^9,37^. Such a comparison would also make it possible to test whether different sensory modalities interact differently with motor-sensory and inter-sensory components in influencing sensorimotor temporal recalibration. Therefore, the impact of sensory modality on recalibration can be tested by sensory events that were not previously associated with the action.

Existing evidence on sensorimotor temporal recalibration suggests that motor-sensory and inter-sensory components are likely modulated by cross-modal interactions. Importantly, studies of modality-specific recalibration have not disentangled how efferent (corresponding to the motor-sensory component) and reafferent (corresponding to the inter-sensory component) feedback contribute to temporal recalibration ^6,9,32,38^. On the other hand, studies investigating the effects of these components have not addressed possible modality-specific effects ^8,22^. To our knowledge, no study has explored the contributions of motor-sensory and inter-sensory components (disentangling efference copy from reafferent feedback) on sensorimotor temporal recalibration while addressing possible cross-modal interactions. Our aim in the present study is to investigate sensorimotor temporal recalibration for motor-auditory and motor-visual events by disentangling the influence of motor-sensory component and inter-sensory component. To this end, we used a recalibration paradigm in which systematic temporal delays between actions and their sensory feedback were introduced (adaptation). To assess temporal recalibration, we tested participants’ perception for variable delays inserted between action-feedback events (test). Crucially, we used active and passive movements at adaptation and test to disentangle motor-sensory and inter-sensory components of recalibration. In the condition where button presses at adaptation and test are both passive (adapt passive, test-passive), a *purely* inter-sensory adaptation (*I* in the above model) between the tactile-proprioceptive and the auditory or visual feedback is expected. Comparing this passive condition with active (voluntary) button presses at adaptation and test (adapt-active, test-active) may aid in understanding the relation between sensorimotor and purely sensory recalibration, and possible recalibration differences as a function of accompanying sensory feedback^6,8,9^. In this condition adaptation is, according to the model, a weighted average of the inter-sensory component *I* and the motor-sensory component *M*. In the third condition, efferent feedback is not present at adaptation, but is present at test (adapt-passive, test-active). Because efferent feedback cannot be adapted, any adaptation effect we observe under this condition should result from the inter-sensory component *I*, but according to the model weighted by its relative reliability. Examining differences between this passive-active and the active-active condition (in which the efferent feedback can be adapted) may hence address the specific contribution of the two components.

Possible mechanisms underlying sensorimotor temporal recalibration involve realignment of either the motor or the sensory event, or both events in time, but also a widening of the temporal window of integration between the events ^39^. These mechanisms are not mutually exclusive, and can all contribute to recalibration ^39^. In the present study, we tested for these possibilities by comparing detection thresholds for delay and just-noticeable differences (JNDs) across delays.

We hypothesized that, if the motor-sensory component is more reliable due to predictions based on efference copy ^16–18,23^, which enhances the temporal organization of events ^12,15,19–21^, then larger adaptation effects should be present for sensorimotor recalibration (adapt-active, test-active condition) than for purely sensory recalibration (adapt-passive, test-passive condition). We further hypothesized that sensorimotor temporal recalibration would result from a combination of motor-sensory components (corresponding to predictions based on efference copy) and inter-sensory components (corresponding to reafferent feedback) weighted according to their reliability ^23,25–27^. If efferent feedback is more reliable than reafferent feedback in terms of recalibrating temporal discrepancies in action-feedback pairs ^8^, we expect lower weighting of inter-sensory discrepancies, and higher weighting of motor-sensory discrepancies. This would suggest that negligible or no recalibration would occur in the adapt-passive, test-active condition compared to the adapt-active, test-active condition. Alternatively, if reafferent feedback is more reliable for recalibration ^22^, then similar adaptation profiles should be evident in the adapt-active, test-active and adapt-passive, test-active conditions, as both conditions contain reafferent feedback (inter-sensory component) in the adaptation phase. Differences across adapted sensory modalities, if found, would point to a change in the relative weighting of the motor-sensory or the inter-sensory component depending on the reliability of the adapted sensory event.

Finally, we hypothesized that a transfer from an adapted to a non-adapted sensory modality would indicate supramodal recalibration, which is not influenced by the sensory feedback of the action ^6,9^, whereas lack of transfer would suggest a modality-specific influence on recalibration ^10,37^. Importantly, the existence of transfer, along with differences in recalibration across adapt-active, test-active and adapt-passive, test-active conditions, would provide insight into which component might receive a higher weight. For example, if the adapted sensory modality is shifted in time, then there should be no cross-modal transfer. Likewise, transfer of recalibration from vision to audition in the adapt-active, test-active but not in the adapt-passive, test-active condition would indicate that the motor-sensory component is shifted in time.

## Methods

### Participants

The experiment was approved by the local ethics committee and was performed in accordance with the Declaration of Helsinki^44^. A total of 14 university students from Philipps University Marburg took part in the study. Data from two participants were discarded from group-level analyses (see Results section), resulting in a final sample of 12 participants (seven females, mean age 24.9±1.64). All participants were right-handed as confirmed by the Edinburgh Handedness Inventory ^45^. They reported normal or corrected-to-normal vision, and normal hearing. In addition, none reported having current psychiatric or neurological conditions or the use of related medication. The participants received monetary compensation for their participation.

### Stimuli and apparatus

Auditory stimuli consisted of brief sine-wave tones (frequency = 2000Hz, duration = ~33.4ms with 2ms rise/fall slopes), and were presented via headphones. Visual stimuli were Gabor patches (2.56°, spatial frequency = 2cycles/degree, duration = ~33.4ms), and were presented on a 24” computer monitor (1920 x 1200pixels resolution, 60Hz frame refresh rate). Stimulus presentation and response recording were controlled by Octave and Psychtoolbox-3 ^46^. Delay detection responses were recorded via a keyboard (‘V’ and ‘N’ buttons on the keyboard).

The experiment was conducted in a dimly lit room. Participants sat at a desk in front of a monitor with a viewing distance of approximately 55cm. Their right index finger was placed on a custom-made button. The custom-made electromagnetic button was used to trigger auditory and visual stimuli. Stroke length of the button was 5mm with a light-barrier triggered within the last 0.2mm of movement. Both voluntary manual and externally activated button presses were recorded by the computer as a left click of a USB mouse. Therefore, jitter and delay of the button press did not depend on whether the movement was active or passive. To ensure that the index finger was pulled by the button for passive movements, cotton bandages were used to fix the finger to the button. The cotton bandages were used during the execution of active movements as well. For active button presses, the initial force was 1.5Newton (N), as measured by a spring force gauge, slowly increasing to approximately 2.5N in the final position. For passive button presses, the finger was initially pulled with approximately 1N, and the force increased to approximately 4N in the final position. The duration of the button press for the passive movement was set to 300ms based on previous studies ^11,49^. For both movements, auditory or visual stimuli were presented only when the button was pressed down completely. In addition, a cushion was provided to ensure a comfortable hand/forearm positioning. The button pad was covered with a black box to prevent the participant from using visual cues from their hand or finger to perform the delay detection task. White noise was presented throughout the experiment to mask any auditory cues, especially the mechanical sound from the passively pressed button. Earplugs were worn by the participants to attenuate any external sound.

### Design

The experimental design involved four within-subjects factors. The first factor corresponded to the *adaptation, test mode*, defining the action to be performed in the adaptation and test phases. The action could either be active in which the participant pressed a button, or passive in which the button was depressed automatically without the participant pressing the button down. Actions in the *adaptation, test mode* pairs could be adapt-active, test-active (sensorimotor recalibration), adapt-passive, test-passive (sensory-sensory recalibration), or adapt-passive, test-active (inter-sensory component in sensorimotor recalibration). The second factor concerned the modality of sensory feedback in the adaptation phase (*adaptation modality*) that could either be auditory (adapt-A) or visual (adapt-V). The third factor corresponded to the modality of sensory feedback during test (*test modality*), that could be either within-modality test (adapt-A, test-A; adapt-V, test-V) or cross-modal test (adapt-A, test-V; adapt-V, test-A). Finally, the fourth factor was the *adaptation delay*, which was the time between the action and the sensory feedback at adaptation, and effects of this factor indicate whether adaptation took place. When *adaptation delay* was 150ms, we expected a decrease in the perceived delay between action and feedback in the test phase, compared to when it was 0ms. The dependent variable was delay detection judgments from six delays (0, 66.8, 133.6, 200.4, 267.2, 334ms) inserted between an active or a passive button press, and auditory or visual feedback at the test phase. A schematic of the experimental conditions is outlined in Figure 1.

**Figure 1.**
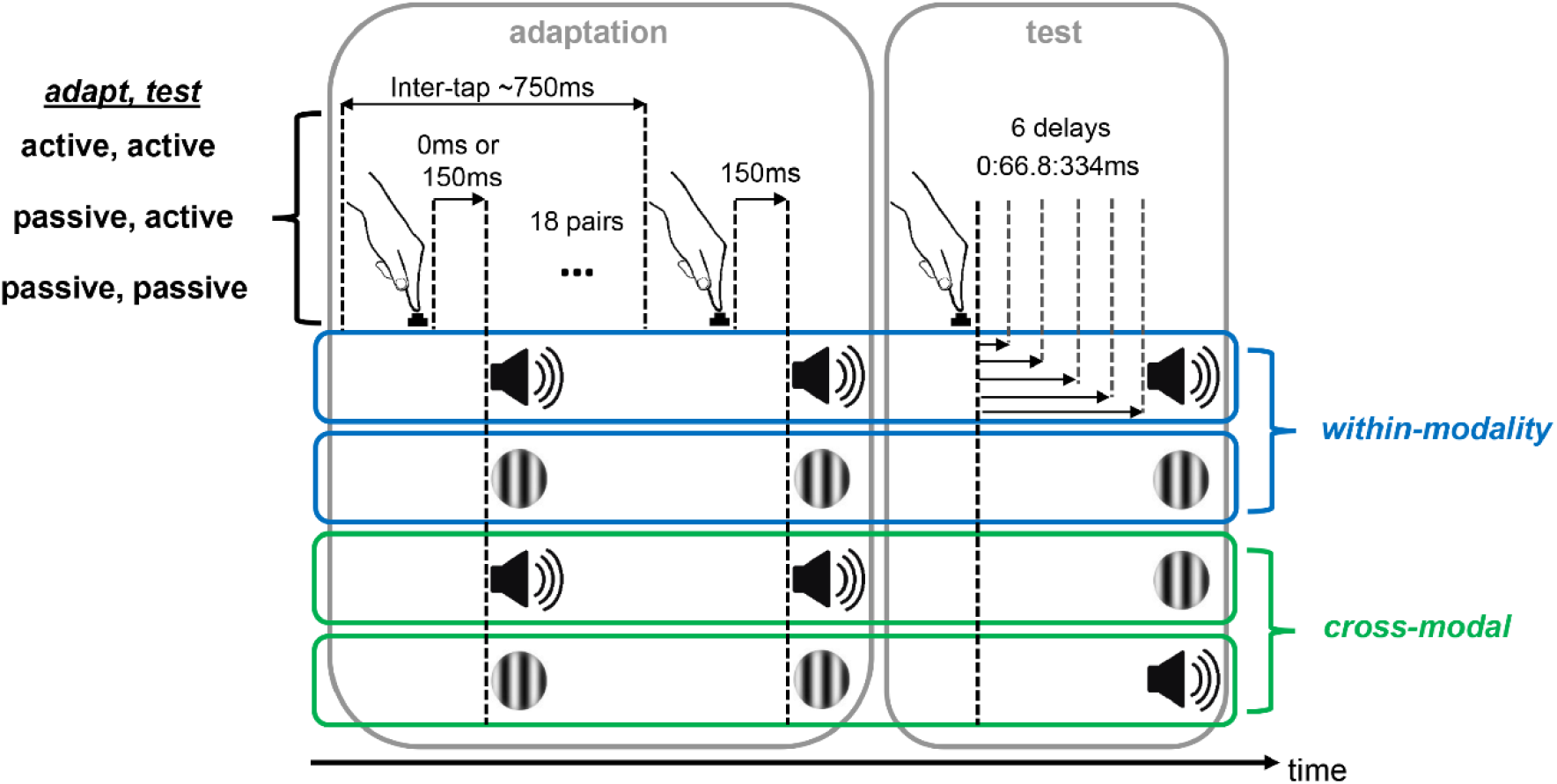
Schematic of the experimental conditions. In the adaptation phase, participants adapted to a systematic delay of either 0ms or 150ms between button press and sensory feedback. Inter-tap interval between the button press and the sensory feedback was ~750ms. In the test phase, the sensory feedback was presented with variable delays (0-334ms), and participants were asked to report whether they detected such a delay.

Participants attended eight experimental sessions completed on separate days. In each session, the participants completed six blocks of different experimental conditions. Adaptation delays were presented on separate days in order to prevent possible carry-over effects ^13^. *Adaptation modality* was kept constant within a specific day, while *adaptation, test mode* and *test modality* were presented on the same day in a pseudorandomized order. We split the experimental conditions in half, so that the first and last four sessions consisted of the same conditions. This was necessary to have a consistent estimate of the detection responses per condition without exhausting the participants.

### Procedure

Each block consisted of 18 trials, all of which involved an adaptation phase and a test phase. On each trial, there were 18 button press-sensory feedback events at adaptation, and six button press-sensory feedback pairs with variable delays at the test phase. The number of repetitions in the adaptation phase and the delays in the test phase were determined based on two previous pilot studies. In the adaptation phase, the participants were asked to perform button presses at a constant pace. Each button press would lead to an auditory or visual stimulus. For trials in which the participant would actively initiate button presses (adapt-active), they were required to press the button approximately every 750ms. For trials in which the button was pressed automatically (adapt-passive), they were asked to let the button go down, and not exert any force on the button. In the test phase, the button presses could either be active (test-active) or passive (test-passive), this time, leading to the presentation of an auditory or a visual stimulus with variable delays. The participants were asked to detect a delay between the button press and the stimulus.

A schematic of an experimental trial is shown in Figure 2. Each block began with instruction of the movement type (button press: active or passive) as well as the modality at adaptation (auditory or visual) for 1500ms. During this time a fixation cross was presented, which remained on the screen for 300ms after the instruction. The fixation cross then disappeared, prompting participants to perform button presses at the instructed pace actively or to let their finger passively press the button. Each button press triggered a beep or the occurrence of a Gabor patch on the screen, either immediately (0ms delay) or delayed in time by 150ms. After the completion of 18 button presses, an instruction followed for 1500ms, informing the participant about movement type and sensory modality in the upcoming test phase (active/passive auditory/visual). The fixation cross disappeared again, prompting the participant to press the button, this time, once. Each button press triggered a beep or a Gabor patch with one of the following six delays: 0, 66.8, 133.6, 200.4, 267.2, and 334ms. A 500ms interval followed in which the fixation cross appeared again. After this interval, the question ‘Delay?’ appeared on the screen. The participants used the keys ‘V’ and ‘N’ for ‘Yes’ and ‘No’ to provide their responses. They had to register their responses within 2000ms. The assignment of keys to yes/no responses was counterbalanced across participants. Six test trials were presented at each test phase, with all trials having one of the six delays. The order of delay presentation was pseudorandomized within a block so that all delays appeared equally often at each position in the trial. The test phase was followed by an inter-trial interval ranging from 500-1500ms. For trials in which the button was actively pressed at adaptation, participants were informed how to adjust their pace of button pressing on the next trial. If the overall (median) interval between button presses fell within the range of 600-900ms, a ‘keep the pace’ instruction appeared for 1500ms after the test phase and prior to the inter-trial interval. If the median intervals were below 600ms and above 900ms, participants received ‘Slower’ and ‘Faster’ instructions, respectively. These *slow* or *fast* trials were immediately repeated until the median button press interval was within the expected range. Overall, most participants were able to keep the pace (2.8% of all trials had to be repeated).

**Figure 2.**
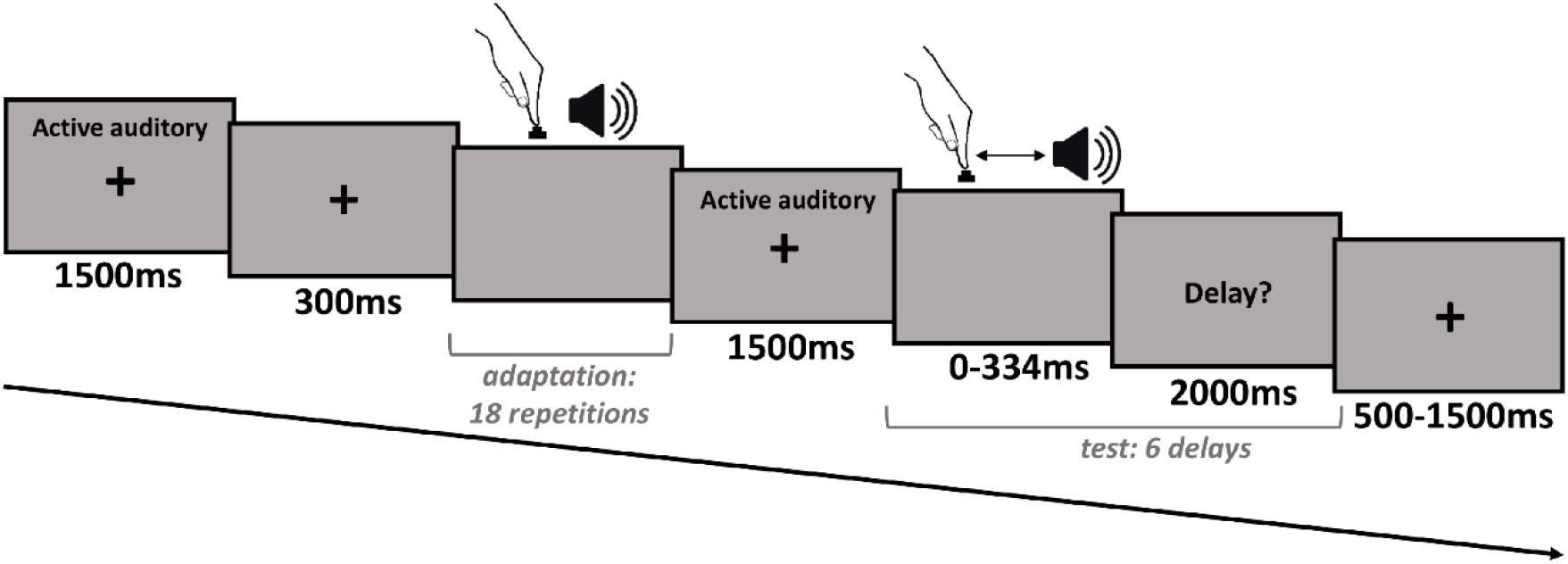
Timeline of an experimental trial. After the disappearance of the fixation cross, participants were instructed to actively initiate 18 button presses each followed by a brief tone. They then received the test phase instruction, in which they were asked to actively press the button again that was accompanied by a tone, and judge whether there was a delay between these two events.

Prior to the experimental blocks, participants practiced tapping for 3 minutes in order to familiarize themselves with the correct pace of the button presses. The participants were also asked whether they felt comfortable with their performance, and if not, were offered further practice. All participants were able to tap in accordance with the auditory signal in the initial training. The tapping practice was followed by short training blocks of the experimental conditions. There were three trials in each training block. The training blocks were presented before the respective experimental blocks. The duration of the entire experiment over the eight sessions was approximately 10.5 hours.

### Data Analysis

The data were subjected to a two-stage inspection procedure. First, we identified extreme detection responses for each participant separately. Second, we checked the overall distribution of detection responses for each condition to determine the appropriate statistical analyses.

In order to discover extreme response patterns, we plotted the proportion of detection responses (i.e., ‘Yes’ responses) as a function of the action-feedback delay at test for each participant and condition. We inspected the detection responses in terms of proportion of detection at the smallest and largest delay trials. For this, we pooled the data across adaptation delays (0 and 150ms) and modalities (within and cross-modal) per recalibration mode. We then calculated the median of proportion detected responses for the largest delay. Data from two participants was excluded from further because their median of proportion detected responses exceeded 50% at no delay trials (suggestive of a strategy to always press ‘yes’, even when there was no delay) or did not exceed 50% at the largest delays (chance-level detection). The remaining data from each participant and condition were fitted to a cumulative Gaussian using psignifit 4.0 ^47^ and Matlab 2019a (Mathworks Inc.). This version of psignifit adopts Bayesian inference to estimate detection parameters ^47^. From fitting psychometric functions, we obtained estimates of detection thresholds (50% point of the psychometric function), and JNDs (difference between the 50% and 84% points of the psychometric function). These estimates were used to conduct statistical analyses. Mauchly’s Test of Sphericity indicated that the assumption of sphericity had not been violated. We performed repeated measures ANOVAs for comparisons across conditions. For each *adaptation, test mode* and *modality* pairing, recalibration was defined as a systematic change in the perception of time as a function of the temporal delay presented at the adaptation phase between the action and the sensory feedback; this would correspond to a change in the detection thresholds or increase in JNDs across delays.

In order to assess the existence of recalibration for different events, and the contribution of motor-sensory and inter-sensory components, we conducted a 3 (*adaptation, test mode*: adapt-active, test-active vs. adapt-passive, test-active vs. adapt-passive, test-passive) x 2 (*adaptation, test modality*: adapt-A, test-A vs. adapt-V, test-V) x 2 (*adaptation delay*: 0ms vs. 150ms) repeated measures ANOVA on thresholds and JNDs.

Results of the first analysis revealed modality-specific effects on sensorimotor temporal recalibration (see Results section). We therefore assessed the existence of cross-modal transfer separately for each adaptation modality. For this, we conducted a 3 (*adaptation, test mode*: adapt-active, test-active vs. adapt-passive, test-active vs. adapt-passive, test-passive) x 2 (*adaptation delay*: 0ms vs. 150ms) analysis on thresholds and JNDs separately for adapt-A, and adapt-V conditions.

Inferential statistics were computed with frequentist hypothesis tests (α = .05). Due to our specific hypothesis regarding adaptation delay (increased thresholds for 150ms compared to 0ms in a specific condition), main effects and post-hoc comparisons were not corrected for multiple comparisons. For other effects, planned post-hoc comparisons were performed when multiple comparisons were made, and Bonferroni correction was applied.

## Results

Table 1 shows group-level threshold and JND estimates for each condition. Figure 3 depicts detection responses as a function of delay for each condition from a representative participant. Overall, detection thresholds were higher in the 150ms condition than in the 0ms condition, indicating recalibration.

**Figure 3.**
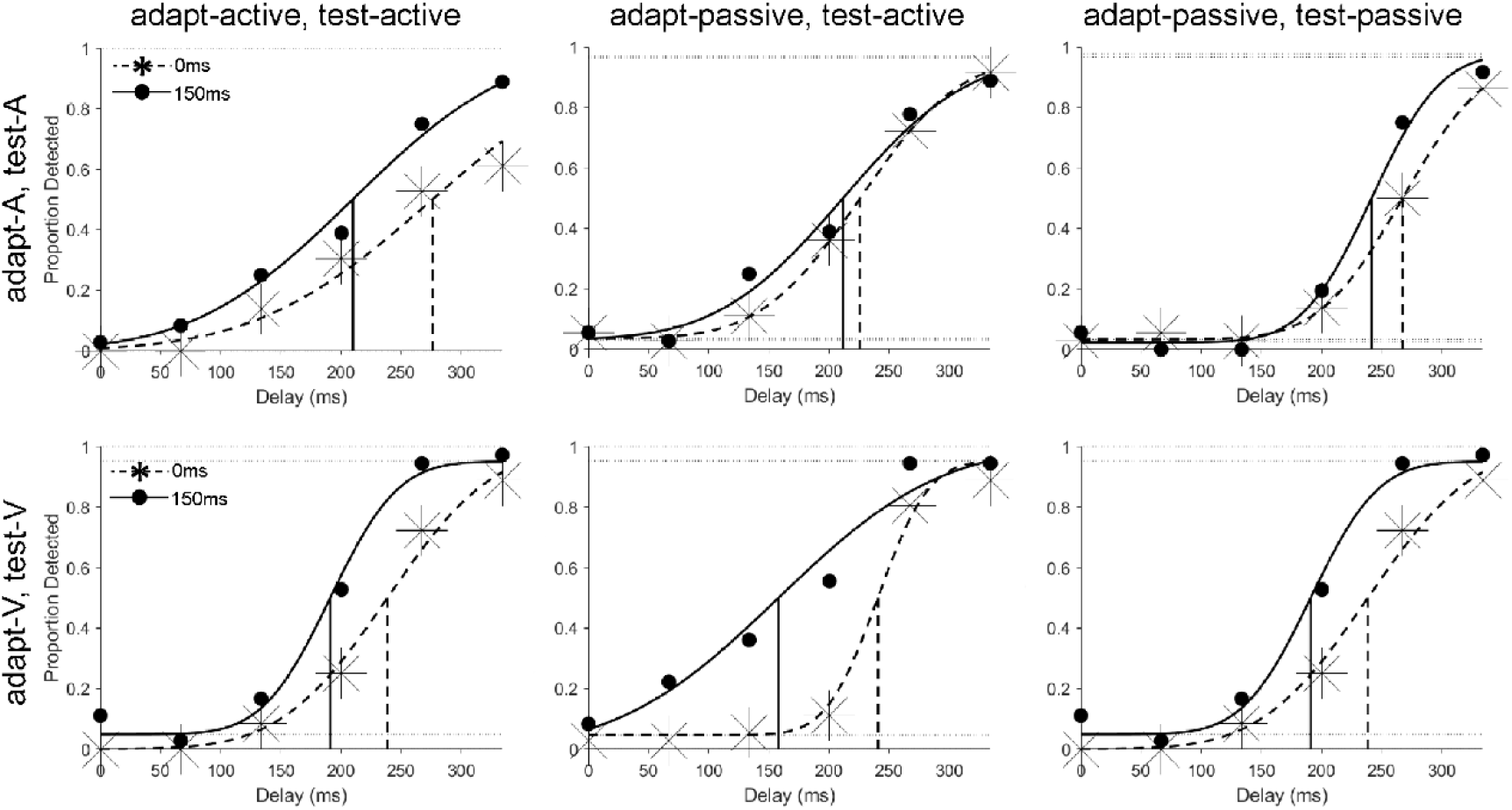
Plots showing detection responses and psychometric function fits for each condition from a representative participant. Filled circles and stars show proportion of detected responses as a function of test delay in the 0ms and 150ms adaptation delay conditions, respectively. A shift in detection thresholds from 0ms to 150ms delay indicates adaptation.

**Table 1.** Mean (standard error of mean, s.e.m.) thresholds and JNDs for each condition.

|  |  | threshold (ms) |  |  |
| --- | --- | --- | --- | --- |
|  |  | adapt-active,<br>test-active | adapt-passive,<br>test-active | adapt-passive,<br>test-passive |
| adapt-A, test-A | 0ms | 202.53 (19.45) | 226.36 (20.25) | 241.83 (13.78) |
|  | 150ms | 229.63 (19.83) | 230.21 (18.02) | 254.44 (11.82) |
| adapt-V, test-V | 0ms | 235.59 (19.98) | 232.87 (21.02) | 236.96 (13.90) |
|  | 150ms | 261.60 (17.88) | 263.82 (18.63) | 258.23 (14.53) |
| adapt-A, test-V | 0ms | 240.06 (21.93) | 243.10 (23.11) | 241.31 (16.84) |
|  | 150ms | 248.32 (16.62) | 252.56 (16.42) | 252.27 (19.76) |
| adapt-V, test-A | 0ms | 225.66 (19.37) | 235.18 (19.07) | 243.57 (20.67) |
|  | 150ms | 248.10 (23.58) | 237.83 (23.23) | 263.55 (15.53) |

|  |  | JND (ms) |  |  |
| --- | --- | --- | --- | --- |
|  |  | adapt-active,<br>test-active | adapt-passive,<br>test-active | adapt-passive,<br>test-passive |
| adapt-A, test-A | 0ms | 75.84 (9.97) | 73.50 (7.07) | 71.02 (10.78) |
|  | 150ms | 65.80 (8.39) | 68.94 (4.81) | 70.67 (12.61) |
| adapt-V, test-V | 0ms | 74.24 (9.44) | 81.82 (13.12) | 72.09 (11.48) |
|  | 150ms | 62.30 (5.05) | 66.53 (5.95) | 67.62 (6.50) |
| adapt-A, test-V | 0ms | 71.06 (8.59) | 72.06 (11.87) | 79.57 (13.20) |
|  | 150ms | 72.75 (11.91) | 80.88 (9.64) | 77.96 (11.49) |
| adapt-V, test-A | 0ms | 68.88 (10.47) | 76.45 (10.75) | 74.19 (9.82) |
|  | 150ms | 79.03 (9.54) | 72.62 (8.75) | 78.58 (11.83) |

### The impact of motor-sensory and inter-sensory components on sensorimotor temporal recalibration

The 3 (*adaptation, test mode*: adapt-active, test-active vs. adapt-passive, test-active vs. adapt-passive, test-passive) x 2 (*adaptation, test modality*: adapt-A, test-A vs. adapt-V, test-V) x 2 *(adaptation delay*: 0ms vs. 150ms) repeated measures ANOVA on thresholds revealed a main effect of *adaptation delay*; *F*(1,11) = 15.50, *p* = .002, □*_p_^2^* = .59. Detection thresholds in the 0ms delay condition (mean = 229.36, s.e.m. = 17.02) were significantly smaller than those in the 150ms delay condition (mean = 249.65, s.e.m. = 15.27). There was also a two-way interaction between *adaptation, test mode* and *adaptation, test modality*; *F*(1,11) = 13.51, *p* < .001, □*_p_^2^* = .55 (cf. Figure 4), and a three-way interaction between *adaptation, test mode; adaptation, test modality* and *adaptation delay*; *F*(1,11) = 3.77, *p* = .04, □*_p_^2^* = .26 (cf. Figure 5). For the two-way interaction, post-hoc comparisons were performed for *adaptation, test modes* in the adapt-A, test-A condition given similar mean thresholds across movements for the adapt-V, test-V conditions. The planned post-hoc comparisons showed that mean detection thresholds (independent of *adaptation delay*) were significantly smaller in the adapt-active, test-active (mean = 216.08, s.e.m. = 19.34) than in the adapt-passive, test-passive (mean = 248.14, s.e.m. = 12.55) condition for the adapt-A, test-A modality; *t*(11) = −4.30, *p* = .002, *d* = 1.44 (see Figure 4). All other effects were non-significant; *adaptation, test mode*, *F*(1,11) =2.72, *p* = .09, □*_p_^2^* = .20; *adaptation, test modality*, *F*(1,11) = 3.58, *p* = .09, □*_p_^2^* = .25; *adaptation, test mode*adaptation delay*, *F*(1,11) = 2.31, *p* = .12, □*_p_^2^* = .17; *adaptation, test modality*adaptation delay*, *F*(1,11) = 1.18, *p* = .30, □*_p_^2^* =.10.

**Figure 4.**
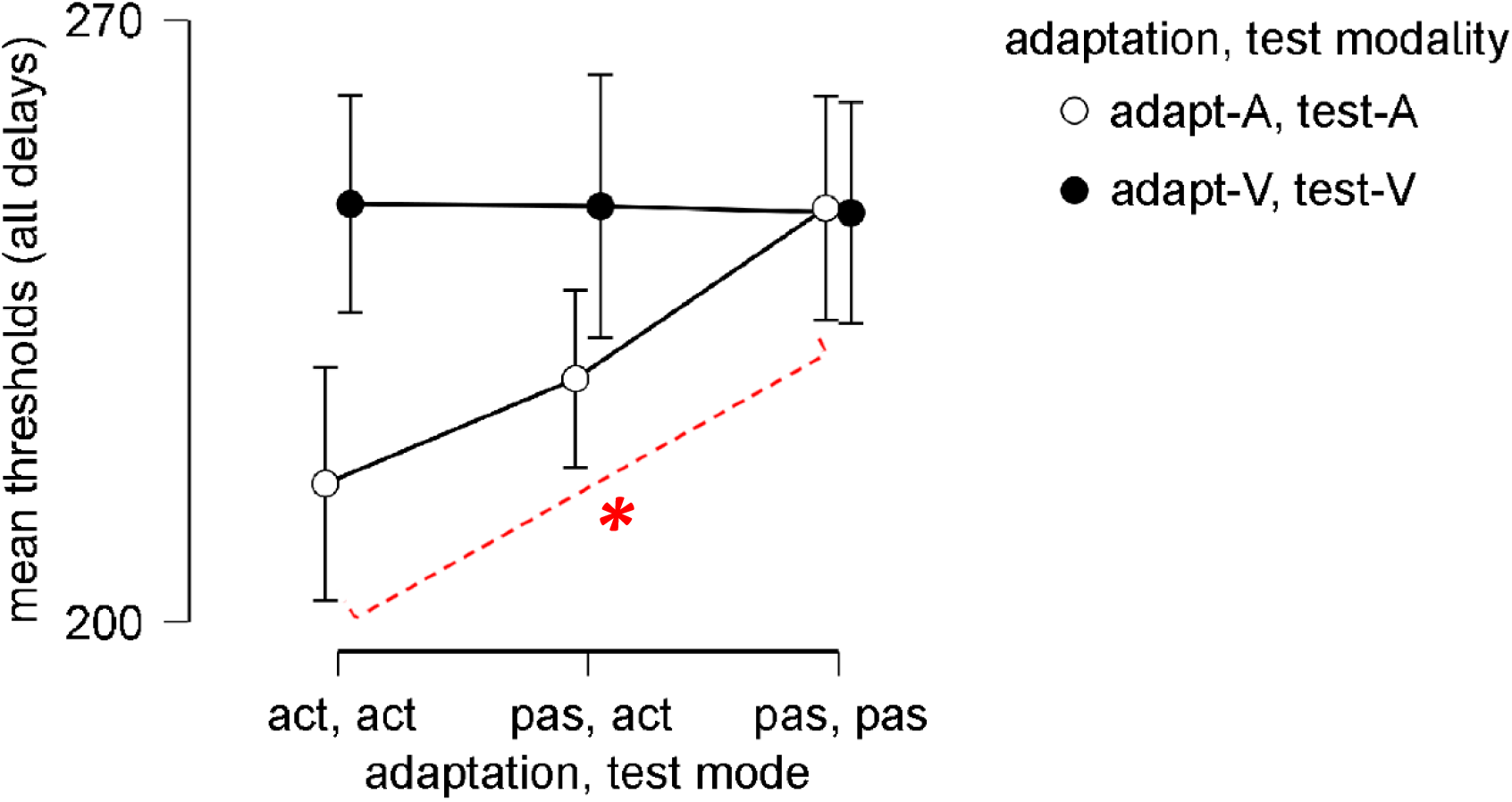
Line plots showing detection thresholds in the adapt-active, test-active (act, act), adapt-passive, test-active (pas, act) and adapt-passive, test-passive (pas, pas) conditions averaged across adaptation delays. Asterisk shows significant differences between conditions. Error bars represent the standard error of the mean (s.e.m.).

**Figure 5.**
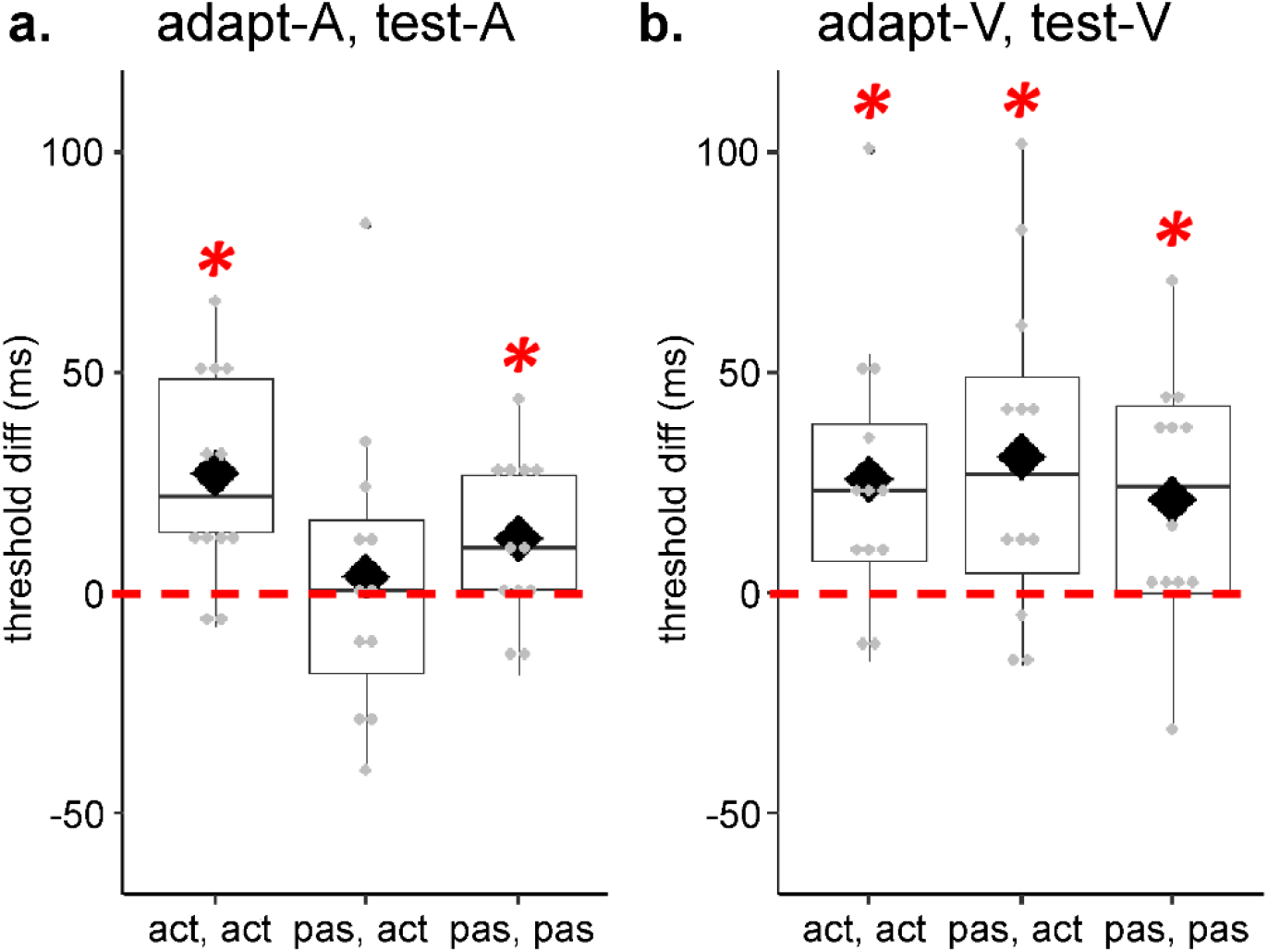
Boxplots with individual data points showing significant effect of adaptation delay (threshold differences between 150ms and 0ms delay conditions) as a function of *adaptation, test mode* in **a.** the adapt-A, test-A condition, and **b.** the adapt-V, test-V condition. The dashed line depicts the difference from 0ms delay. Diamonds and solid lines show the mean and the median values of the data, respectively. Asterisks show significant effects.

For the three-way interaction, we conducted planned comparisons testing the presence of adaptation in each condition, namely the difference between the adaptation delays for each *adaptation, test mode* and *adaptation, test modality*. For each participant, we subtracted the detection thresholds in the 150ms delay condition from the 0ms delay condition separately for *adaptation, test mode* and *modality*, and tested the differences (threshold_Diff_) against 0 with one-sample t-tests. As the expected direction of the effect was specific (larger detection thresholds for the 150ms delay compared to 0ms delay), we used one-tailed t-tests. Moreover, we did not correct for multiple comparisons as the presence of adaptation in each condition was not expected to be dependent on adaptation in another condition. The results showed significantly different threshold_Diff_ values for the following conditions: active-adapt-A, active-test-A; *t*(11) = 3.96, *p* = .001, *d* = 1.14, active-adapt-V, active-test-V; *t*(11) = 2.88, *p* = .007, *d* = .83, passive-adapt-V, active-test-V; *t*(11) = 2.85, *p* = .008, *d* = .82, passive-adapt-A, passive-test-A, *t*(11) = 2.33, *p* = .02, *d* = .67, passive-adapt-V, passive-test-V, *t*(11) = 2.62, *p* = .01, *d* = .76 (see Figure 5).

The 3 (*adaptation, test mode*: adapt-active, test-active vs. adapt-passive, test-active vs. adapt-passive, test-passive) x 2 (*adaptation, test modality*: adapt-A, test-A vs. adapt-V, test-V) x 2 (*adaptation delay*: 0ms vs. 150ms) repeated measures ANOVA on JNDs resulted in no significant differences across conditions. Together, the results suggest a shift in the detection thresholds as a function of adaptation delay for the visual modality independent of action at adaptation or test. For the auditory modality, recalibration is present in the adapt-active, test-active and adapt-passive, test-passive conditions. JNDs across adaptation delays were similar, suggesting no differences in the temporal window of integration for event pairs as a function of delay at adaptation.

### Cross-modal transfer of recalibration

Results of the initial analysis suggest a shift in the detection thresholds with adaptation delays (confirming recalibration), and differences in recalibration across sensory modalities. We therefore examined transfer of recalibration separately for each *adaptation, test modality*. For adapt-A, the 3 (*adaptation, test mode*: adapt-active, test-active vs. adapt-passive, test-active vs. adapt-passive, test-passive) x 2 (*adaptation delay*: 0ms vs. 150ms) repeated measures ANOVA on thresholds for adapt-A, test-V revealed no significant main or interaction effects across conditions; F(1,11) = .44, p = .65, □*_p_^2^* = .04 for *adaptation, test mode*, *F*(1,11) = 2.73, *p* = .13, □*_p_^2^* = .20 for *adaptation delay*, *F*(1,11) = .02, *p* = .98, □*_p_^2^* = .002 for interaction between *adaptation, test mode* and *adaptation delay*.

The 3 (*adaptation, test mode*: adapt-active, test-active vs. adapt-passive, test-active vs. adapt-passive, test-passive) x 2 (*adaptation delay*: 0ms vs. 150ms) repeated measures ANOVA on JNDs for adapt-A, test-V revealed no significant main or interaction effects; *F*(1,11) = 1.38, *p* = .27, □*_p_^2^* = .11 for *adaptation, test mode*, *F*(1,11) = 1.16, *p* = .31, □*_p_^2^* = .10 for *adaptation delay*, *F*(1,11) = .60, *p* = .56, □*_p_^2^* = .05 for interaction between *adaptation, test mode* and *adaptation delay* (see Figure 6a).

**Figure 6.**
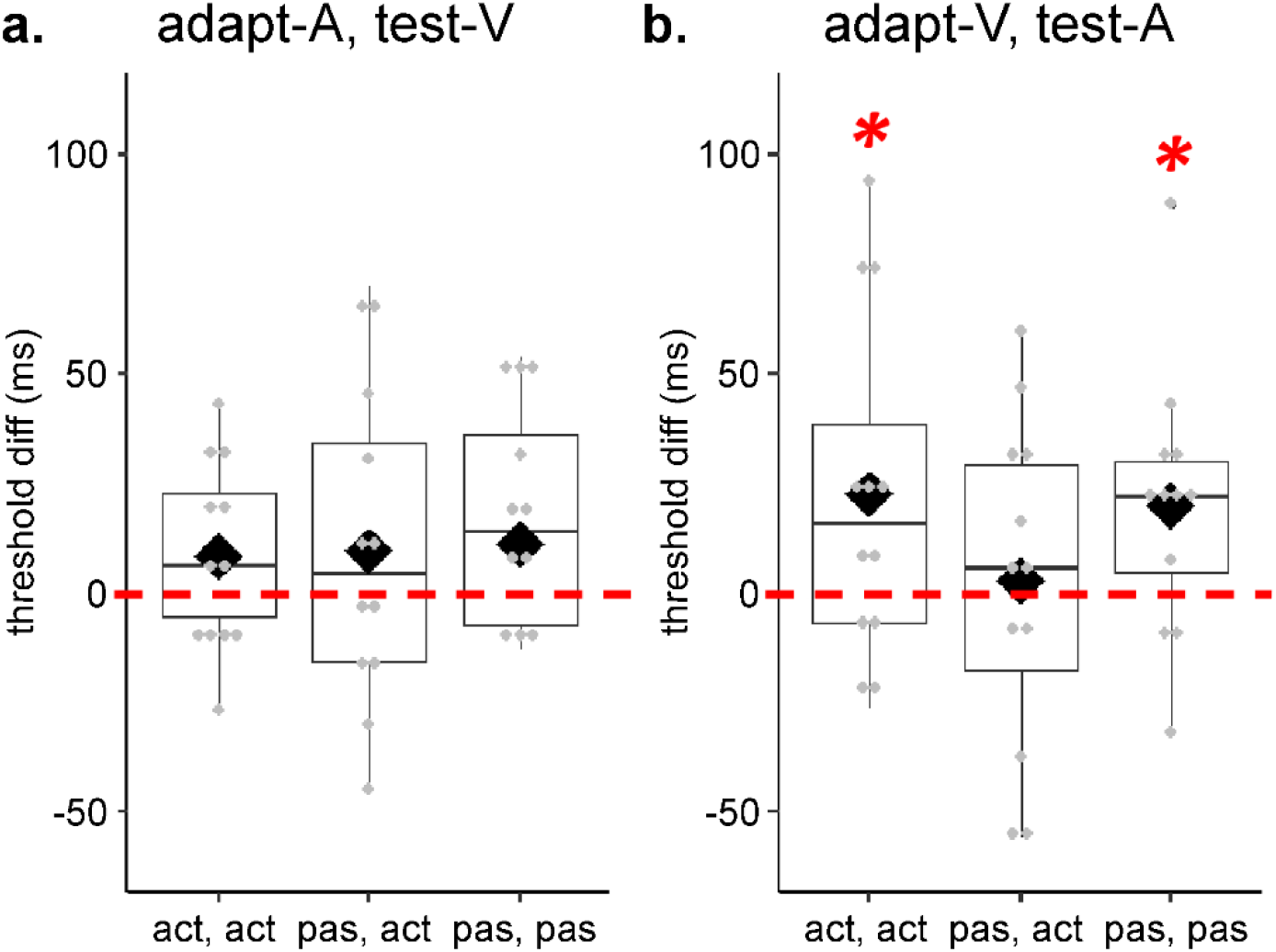
Boxplots with individual data points showing significant effect of adaptation delay (threshold differences between the 150ms and 0ms conditions) **a.** for adapt-A, test-V, and **b.** for adapt-A, test-V across conditions. The dashed line depicts the difference from 0ms delay. Diamonds and solid lines show the mean and the median of the points, respectively. Bold asterisks indicate significant effects.

The 3 (*adaptation, test mode*: adapt-active, test-active vs. adapt-passive, test-active vs. adapt-passive, test-passive) x 2 (*adaptation delay*: 0ms vs. 150ms) repeated measures ANOVA on thresholds for adapt-V, test-A resulted in a main effect of *adaptation delay*; *F*(1,11) = 5.24, *p* = .04, □*_p_^2^* = .32. Other effects were not significant; *F*(1,11) = 2.96, *p* = .07, □*_p_^2^* = .21 for *adaptation, test mode*, *F*(1,11) = 1.23, *p* = .31, □*_p_^2^* = .10 for interaction between *adaptation, test mode* and *adaptation delay*. Although we found a main effect of *adaptation delay* on detection thresholds, indicating cross-modal transfer of recalibration independent of *adaptation, test mode*, an inspection of threshold differences (see Figure 6b) suggests that the effect was mainly driven by the adapt-active, test-active and adapt-passive, test-passive conditions. Indeed, one sample t-tests (one-sided) on the threshold differences for each *adaptation, test mode* confirms this observation; *t*(11) = 2.0, *p* = .04, *d* = .58 for adapt-active, test-active, *t*(11) = .24, *p* = .41, *d* = .07 for adapt-passive, test-active, *t*(11) = 2.27, *p* = .02, *d* = .66 for adapt-passive, test-passive condition (Figure 6b).

The 3 (*adaptation, test mode*: adapt-active, test-active vs. adapt-passive, test-active vs. adapt-passive, test-passive) x 2 (*adaptation delay*: 0ms vs. 150ms) repeated measures ANOVA on JNDs revealed no significant main or interaction effects; *F*(1,11) = .34, *p* = .72, □*_p_^2^* = .03 for *adaptation, test mode*, *F*(1,11) = .57, *p* = .47, □*_p_^2^* = .05 for *adaptation delay*, *F*(1,11) = 1.70, *p* = .21, □*_p_^2^* = .13 for interaction between *adaptation, test mode* and *adaptation delay*.

## Discussion

In the present study, we aimed to disentangle motor-sensory and inter-sensory components of sensorimotor temporal recalibration for visual and auditory feedback. To this end, we presented participants with systematic delays between button presses and sensory feedback, and tested whether detection of variable delays inserted between action-feedback events changed after adaptation. To investigate the role of motor-sensory and inter-sensory components, we used active and passive movements at the adaptation and test phases. We hypothesized that if predictions based on efference copy play a substantial role in adaptation due to their reliability ^8,16–18,23,25,26^, we would observe limited recalibration when the motor-sensory component was absent (equation (1)). Our results indicate that the motor-sensory component determines the temporal recalibration of motor-auditory events, while the inter-sensory component determines the temporal recalibration of motor-visual events. Transfer of recalibration from the visual to the auditory domain, and not from the auditory to the visual domain, highlights the influence of supramodal mechanisms on the inter-sensory component. Our results suggest that the extent to which motor-sensory and inter-sensory components contribute to sensorimotor temporal recalibration depends on the modality of sensory feedback.

Based on previous work on optimal cue combination ^23,25–28,30^, we suggested a model for sensorimotor temporal recalibration which combines motor-sensory (*M*) and inter-sensory (*I*) components in remapping time point estimates of actions and sensory feedback. Our model posits that the reliability of the sensory feedback modality modulates the weight of each component. Overall, shifts in the detection thresholds from 0ms to 150ms delay are smaller for the adapt-passive, test-active condition than for the adapt-passive, test-passive condition. This indicates that recalibration in the adapt-passive, test-active condition reflects only a weighted part of the inter-sensory component, and supports the interpretation of the data in line with the model. Nevertheless, note that this pattern is slightly violated in the adapt-V, test-V condition. However, the similarities in adaptation between the adapt-passive, test-passive and the adapt-passive, test-active condition for visual feedback might be explained by assuming that no weight is given to the motor-sensory component. Apart from threshold differences, we did not find a main effect of adaptation delay on JNDs, suggesting similar temporal integration windows across adaptation delays. However, lack of JND differences across adaptation delays does not necessarily mean that other processes such as temporal window of integration does not contribute to sensorimotor temporal recalibration.

Comparing adaptation across sensorimotor (adapt-active, test-active) and purely sensory (adapt-passive, test-passive) events allowed us to address whether predictions based on efference copy provide additional advantage in recalibrating the timing of related events ^12,15,19–21^, Our analyses of within-modality conditions demonstrate temporal recalibration for sensorimotor as well as purely sensory events, both for auditory and visual feedback. This suggests that the motor-sensory component is not necessarily required in recalibrating the timing of related events. Nevertheless, effect size estimates for auditory recalibration were larger in the adapt-active, test-active condition than in the adapt-passive, test-passive condition. In addition, overall detection thresholds for motor-auditory events in the adapt-active, test-active condition were significantly smaller than detection thresholds in the adapt-passive, test-passive condition, indicating higher sensitivity to discrepancies for actions with auditory feedback. We did not find a difference in detection thresholds between adapt-active, test-active and adapt-passive, test-passive conditions when the sensory feedback was visual. These results align with existing evidence showing improvement of temporal sensitivity between tactile and auditory events when the tactile event is coupled with an action ^43^, suggesting that, at least in the auditory domain, predictions based on efference copy facilitate grouping of events ^12,15,19–21^. However, they do not indicate an overall advantage in recalibrating the perceiving timing of sensorimotor events over purely sensory events.

Apart from differences between sensorimotor and purely sensory events, we addressed the impact of motor-sensory components (corresponding to predictions based on efference copy) and inter-sensory components (corresponding to reafferent feedback) on sensorimotor temporal recalibration. By introducing a condition with passive button presses at adaptation and active button presses at test, we were able to disentangle the impact of efferent from reafferent feedback, both of which may contribute to recalibration ^9,22^ (*M* and *I* in equation (1), respectively). Despite our finding that sensorimotor recalibration for actions with auditory and visual feedback are similar, our results point to enhancement of adaptation when predictions are based on the efference copy, specifically when the adapted modality is auditory. This is evident from the systematic shift in detection thresholds observed for adapt-A, test-A trials in the adapt-active, test-active condition, but not in the adapt-passive, test-active condition ^12,15,19–21^. Together with the finding that temporal sensitivity is in general higher for motor-auditory than for sensory-auditory events, this suggests a specific temporal recalibration of motor-auditory events. What might explain the dependencies between motor and auditory events observed in our study? In the temporal domain, audition is more reliable than other sensory modalities ^33,34^. Moreover, the occurrence of audio-tactile events is susceptible to temporal discrepancies ^48,49^, presumably because in the real world such events frequently occur close in proximity^36^. This might explain the lack of temporal recalibration between button presses and auditory feedback when only the inter-sensory component was available, and efference copy information was absent. Previous work demonstrates that the motor system is more closely linked to the auditory system than to the other senses ^50–53^, making the motor-sensory component dominant for the temporal remapping of actions and auditory events in time. Together, our results support previous findings showing asymmetric transfer of recalibration for sensorimotor events; that is, transfer of recalibration from visual to the auditory domain, but not vice versa^10,37,38^.

Despite the clear advantage of predictions based on efference copy in temporal recalibration for motor-auditory events, a different pattern emerged for motor-visual events. Recalibration was present for event pairs involving visual modality *independent of* whether predictions based on efference copy were present at adaptation. More specifically, adaptation was demonstrated for adapt-active, test-active as well as for adapt-passive, test-active conditions when the sensory event was visual. These results suggest that, compared to the inter-sensory component, the motor-sensory component is assigned higher weight in recalibrating motor-auditory events, while the opposite is true for motor-visual events. This indicates differential contributions of motor-sensory and inter-sensory components on sensorimotor temporal recalibration based on the sensory feedback modality^8,22^. Our results indicate that temporal adaptation for motor-visual events involves remapping of timing between the motor and the visual event driven by the inter-sensory component; that is, the adaptation mainly occurs between tactile-proprioceptive feedback from the action and visual feedback of the action ^22,23,25^.

In order to better explain the mechanism underlying sensorimotor temporal recalibration, we assessed the influence of cross-modal transfer of recalibration. Transfer of recalibration from one modality to another would favor a supramodal mechanism in driving recalibration, whereas lack of transfer would indicate modality-specific effects on recalibration ^6,9,10,37^. In other words, if sensory feedback of the action does *not* influence recalibration, then we should observe transfer of recalibration across modalities. Our results do not indicate a transfer from audition to vision. Note that this finding is independent of whether the adapted event involved an action or not. On the other hand, we found transfer of adaptation from vision to audition for sensorimotor and sensory-sensory events. The transfer from the visual to the auditory domain points to supramodal recalibration that is *not* driven by the sensory feedback. Together with the finding that temporal recalibration for visual events was independent of the motor-sensory component, this suggests supramodal recalibration of sensorimotor events based on the inter-sensory component. Nevertheless, lack of transfer from the visual to the auditory modality when efference copy was absent during adaptation (adapt-passive, test-active condition) indicates that the motor-sensory component still contributes to sensorimotor recalibration.

To conclude, our results demonstrate that sensorimotor temporal recalibration results from interactions between motor-sensory and inter-sensory components. These interactions are not only modulated by the efference copy and reafferent feedback, but also by the sensory feedback modality associated with the action. We suggest that incongruent results on sensorimotor temporal recalibration within and across different sensory modalities can be explained by the differential contributions of motor-sensory and inter-sensory components, which might further depend on the reliability of the adapted sensory event.

## Acknowledgements

We thank Jacob R. Cheeseman for his valuable editorial assistance with the manuscript, and Philipp Lange for help with data collection. This study is funded by the Deutsche Forschungsgemeinschaft (DFG, German Research Foundation) – project number 222641018 – SFB/TRR 135 TP A3. Benjamin Straube is funded by the DFG (STR 1146/15-1 grant number 429442932, STR 1146/9-1/2, grant number 286893149).

## CreDit Authorship Contribution Statement

BEA: Conceptualization, methodology, investigation, software, data curation, formal analysis, visualization, writing – original draft, reviewing and editing. BvK: Conceptualization, methodology, writing – reviewing and editing. KF: Writing – reviewing and editing. TK: Conceptualization, writing – reviewing and editing, funding acquisition. KD: Conceptualization, methodology, formal analysis, writing – reviewing and editing, supervision. BS: Conceptualization, methodology, formal analysis, writing – reviewing and editing, funding acquisition, supervision, project administration.

## Additional Information

The authors declare no competing financial interests.

The data will be provided online at the pre-registered doi: osf.io/c27×6/, and can be used for noncommercial research purposes upon acceptance of this article for publication.

